# Domains of transmission and association of community, school, and household sanitation with soil-transmitted helminth infections among children in coastal Kenya

**DOI:** 10.1101/649509

**Authors:** William E Oswald, Katherine E Halliday, Carlos Mcharo, Stefan Witek-McManus, Stella Kepha, Paul M Gichuki, Jorge Cano, Karla Diaz-Ordaz, Elizabeth Allen, Charles S Mwandawiro, Roy M Anderson, Simon J Brooker, Rachel L Pullan, Sammy M Njenga

## Abstract

**Introduction:** Few studies have simultaneously examined the role of sanitation conditions at the home, school, and community on soil-transmitted helminth (STH) infection. We examined the contribution of each domain that children inhabit (home, village, and school) and estimated the association of sanitation in each domain with STH infection.

**Methods:** Using data from 4,104 children from Kwale County, Kenya, who reported attending school, we used logistic regression models with cross-classified random effects to calculate measures of general contextual effects and estimate associations of village, school, and household sanitation with STH infection.

**Findings:** We found reported use of a sanitation facility by households was associated with reduced prevalence of hookworm infection but not with reduced prevalence of *T. trichiura* infection. School sanitation coverage > 3 toilets per 100 pupils was associated with lower prevalence of hookworm infection. School sanitation was not associated with *T. trichiura* infection. Village sanitation coverage > 81% was associated with reduced prevalence of *T. trichiura* infection, but no protective association was detected for hookworm infection. General contextual effects represented by residual heterogeneity between village and school domains had comparable impact upon likelihood of hookworm and *T. trichiura* infection as sanitation coverage in either of these domains.

**Conclusion:** Findings support the importance of providing good sanitation facilities to support mass drug administration in reducing the burden of STH infection in children.

**Author Summary:** Infection by whipworm and hookworm results from either ingestion of eggs or larvae or through skin exposure to larvae. These eggs and larvae develop in suitable soils contaminated with openly-deposited human faeces. Safe disposal of faeces should reduce transmission of these soil-transmitted helminths (STH), yet evidence of the impact of sanitation on STH transmission remains limited. We used data collected during a large, community-wide survey to measure prevalence of STH infections in coastal Kenya in 2015 to examine the relationship between sanitation conditions at home, school, and village and the presence of STH infection among 4,104 children who reported attending schools. We found that sanitation access at home and school sanitation coverage, but not the overall level of village sanitation coverage, was protective against hookworm infection. In contrast, only high village sanitation coverage, but not home or school sanitation, was protective against whipworm infection. Current STH control strategies emphasise periodic deworming through mass drug administration (MDA) of at-risk populations, including school-age children. Our findings highlight the need for continued efforts, alongside MDA, to extend access to good sanitation facilities at homes, schools, and across communities.

## Introduction

In 2016, it was estimated that world-wide more than 1.5 billion people were infected with at least one species of soil-transmitted helminth (STH)(1). Infection by roundworm, *Ascaris lumbricoides*, and whipworm, *Trichuris trichiura*, results from ingestion of embryonated eggs, and the hookworms, *Necator americanus* and *Ancylostoma duodenale*, infect humans through skin exposure to or ingestion of larvae (*A. duodenale*) that develop in warm, moist soils from eggs in openly-deposited human faeces (2, 3). Preventing human contact with excreta through consistent safe disposal of faeces should reduce STH transmission, yet evidence of the impact of sanitation on STH remains limited (4–6). The concept of private and public domains of transmission for STH has been described previously (7, 8), but to our knowledge few studies have simultaneously examined the role of sanitation conditions at the home, school, and community on STH infection (9–11).

Multilevel statistical models provide a means to estimate effects of individual factors, using measures of association. They can simultaneously assess general contextual effects upon individual health outcomes, using measures of within-unit clustering and between-unit heterogeneity (12, 13). Such models provide a useful complement to mathematical modelling of transmission dynamics for examining specific effects and possible areas for intervention (14). Identifying effects of sanitation within each of the domains that a child inhabits will contribute to evidence for the prioritization of sanitation promotion activities alongside the regular deworming of at-risk populations, including school-aged children, recommended by the World Health Organization (15, 16).

Employing multilevel modelling, we investigate the relative importance of household, village and school domains and estimate the association of sanitation in each domain on STH infection among Kenyan children who attend school.

## Methods

### Study Population

The study took place in Kwale County on the south Kenyan coast. Data were collected during a cross-sectional parasitological survey conducted between March-May 2015 as the baseline for TUMIKIA, a randomised, controlled trial to evaluate the impact and cost-effectiveness of school-based versus community-wide deworming on STH transmission (NCT02397772, www.clinicaltrials.gov).

Study design, baseline findings, and impact have been described previously (17, 18) (Halliday KE, Oswald WE, Mcharo C, Beaumont E, Gichuki PM, Kepha S, et al. Community-level epidemiology of soil-transmitted helminths in the context of school-based deworming: Baseline results of a cluster randomised trial on the coast of Kenya. In Press). For the baseline survey, 225 households were randomly selected within 120 community units (CUs) of approximately 1000 households comprising 2 to 7 villages. Among consenting households, a structured questionnaire was conducted with the head of household or primary caregiver to collect information on demographics, ownership of key assets, and sanitation, hygiene, and water conditions. One household member (aged ≥ 2 years) was randomly selected to provide a stool sample. A questionnaire was then conducted with individuals who provided samples or their caregiver to collect information on deworming within the last year and observe their footwear. School facility surveys were conducted across Kwale County in June 2015 and July 2016. During visits, student enrolment was recorded from school registers, and sanitation conditions were assessed using structured observations. All data were collected on smartphones running the Android operating system (Google, Mountain View, CA, USA) using SurveyCTO (Dobility, Inc., Cambridge, MA, USA). Records from school and household surveys were linked based on the school each child reported attending. Geographic coordinates (based on WGS84 datum) were systematically collected at each household and school using the smartphones’ global positioning systems. Missing coordinates for 4 schools were obtained from Google Maps (Google, Mountain View, CA, USA). Children were excluded *a priori* if they lived in villages or attended schools in semi-arid areas unsuitable for STH transmission (Halliday KE, Oswald WE, Mcharo C, Beaumont E, Gichuki PM, Kepha S, et al. Community-level epidemiology of soil-transmitted helminths in the context of school-based deworming: Baseline results of a cluster randomised trial on the coast of Kenya. In Press). Children were eligible if they were sampled in the 2015 TUMIKIA baseline parasitological survey, aged 5 to 14 years, and reported attending school.

### STH infection

Kato-Katz microscopy was used to enumerate STH eggs (*A. lumbricoides, T. trichiura*, and hookworm) per gram of stool. For both hookworm and *T. trichiura*, our outcome was a dichotomous indicator for the presence of > 0 eggs in stool samples (i.e. prevalence). *A. lumbricoides* was not examined in detail since few cases were detected. STH infection was also classified based on categories of infection intensity (19), and frequencies were tabulated across categories of household, community, and school sanitation as detailed below.

### Sanitation measures

The measure for household sanitation access was combined from reported use of a toilet facility on or off the household’s compound. Using all households sampled per village for the TUMIKIA baseline survey, this measure of household sanitation was aggregated for village sanitation coverage, as the percentage of households with reported access to sanitation. During structured observations at schools, the number of latrines considered usable (not assigned to teachers, locked, or with full pits) was quantified, excluding urinals, for both girls and boys. School sanitation coverage was calculated as the number of usable toilets per enrolled pupil, in contrast to the indicator of students per toilet (20). Village and school sanitation coverage were categorised based on estimated quartiles to explore possible non-linear relationships during modelling.

### Covariates

Information on covariate specification and creation of household socioeconomic status measure is described elsewhere (Halliday KE, Oswald WE, Mcharo C, Beaumont E, Gichuki PM, Kepha S, et al. Community-level epidemiology of soil-transmitted helminths in the context of school-based deworming: Baseline results of a cluster randomised trial on the coast of Kenya. In Press). One school surveyed in 2015 was missing total enrolment, so 2016 enrolment was used. For 51 schools attended by children for which we had no 2015 data, enrolment and sanitation conditions from the 2016 survey were used. Environmental and sociodemographic conditions hypothesized to influence STH transmission and sanitation coverage were assembled in a geographic information system using ArcGIS 10.5 (ESRI, Redlands, CA, USA), and values were extracted for each school and household (Halliday KE, Oswald WE, Mcharo C, Beaumont E, Gichuki PM, Kepha S, et al. Community-level epidemiology of soil-transmitted helminths in the context of school-based deworming: Baseline results of a cluster randomised trial on the coast of Kenya. In Press). We aggregated mean continuous and mode categorical values per village, using all households sampled per village.

### Ethical Approval

The TUMIKIA trial protocol was approved by the Kenya Medical Research Institute and National Ethics Review Committee (SSC Number 2826) and the London School of Hygiene & Tropical Medicine (LSHTM) Ethics Committee (7177). Written informed consent was sought from the household head or adult answering the household-level questionnaire and from the individual selected to provide the stool sample and complete the individual-level questionnaire. Parental consent was sought for individuals aged 2 to 17 years and written assent was additionally obtained from children aged 13 to 17 years. All information and consent procedures were conducted in Kiswahili.

### Statistical analyses

We estimated associations between sanitation conditions in the domains of interest and presence of hookworm and *T. trichiura* infection, separately, using logistic regression models with cross-classified (non-nested) random effects to account for membership of children within village of residence and school attended.

We fit a series of generalised mixed models (with logit link), excluding observations with missing outcome or covariate information, which assumes the probability of having complete data is independent of the outcome after adjusting for included covariates. First, we fit an intercept-only model containing school- and village-specific random effects to quantify between school and between village variation in STH infection. Next, we fit models with fixed effects for sanitation conditions in each domain separately and then together in a combined unadjusted model. Finally, we fit a model containing all sanitation effects, adjusting for potential confounders. Confounders were selected based on existing knowledge and encoding possible causal relationships in directed acyclic graphs. We then implemented d-separation in DAGitty to identify minimal sufficient sets of available covariates to control to estimate effects of sanitation on both hookworm (S2 Supporting Information) and *T. trichiura* infections (S3 Supporting Information)(21, 22). Estimation used Hamiltonian Monte Carlo sampling to improve mixing and reduce auto-correlation (23). We conducted a sensitivity analysis excluding outliers in and missingness of distance from child’s home to reported school attended.

We report prevalence odds ratios (PORs) with 95% credible intervals (CIs) as fixed effects for sanitation in each domain. We also calculated measures to quantify general contextual effects on individual infection (13). Proportion of total observed individual variation in the outcome attributable to between-school and between-village variation, or intraclass correlation coefficient (ICC), was calculated using the latent variable method. This method converts individual variance to the logistic scale from the probability scale and assumes an underlying continuous propensity for infection following a logistic distribution with individual variance of π^2^/3 (24). We calculated median odds ratios (MORs), as measures of residual heterogeneity on the odd ratio scale. MORs are always greater than or equal to 1 (MOR = 1 if there is no variation between areas) and interpreted as the median value of the odds ratios between comparable individuals drawn randomly from high and low risk areas. MORs are useful as they measure how much individual infection is determined by domain membership and are directly comparable to fixed effects (24). The ICC is recommended, however, for estimating general contextual effects as a measure of clustering that incorporates both between- and within-area variance (12). We calculated 80% interval odds ratios (80% IORs) for village and school sanitation fixed effects. This measure does not reflect the estimate’s precision but instead is recommended to consider residual variation in the interpretation of fixed effects (25). A wider interval indicates greater unexplained between-area variation, and the inclusion of 1 indicates between-area variance is large compared to the specific fixed effect. Analyses were conducted in Stata 15 (StataCorp LP, College Station, TX, USA) and in R 3.5 (r-project.org) using the ‘brms’ package.

## Results

Of 6,066 school-aged children with matched samples in the survey, 602 (10%) reported not attending school, and 688 (11%) reported attending school but resided in a village or were linked to a school in a semi-arid area (Figure 1). Among these excluded children, prevalence of infection was 3% (22/688), < 1% (4/688), and 0% (0/688) for hookworm, *T. trichiura* and *A. lumbricoides*, respectively. Of 4,776 eligible children, the school reported to be attended was identified for 4,243 children (89%) and school survey data was available for 4,163 children (87%). Eligible children without school information were younger, more often being boys, less likely to have been dewormed, and less poor (S4 Table). Data were available for 4,104 eligible children (86%).

**Figure 1.**
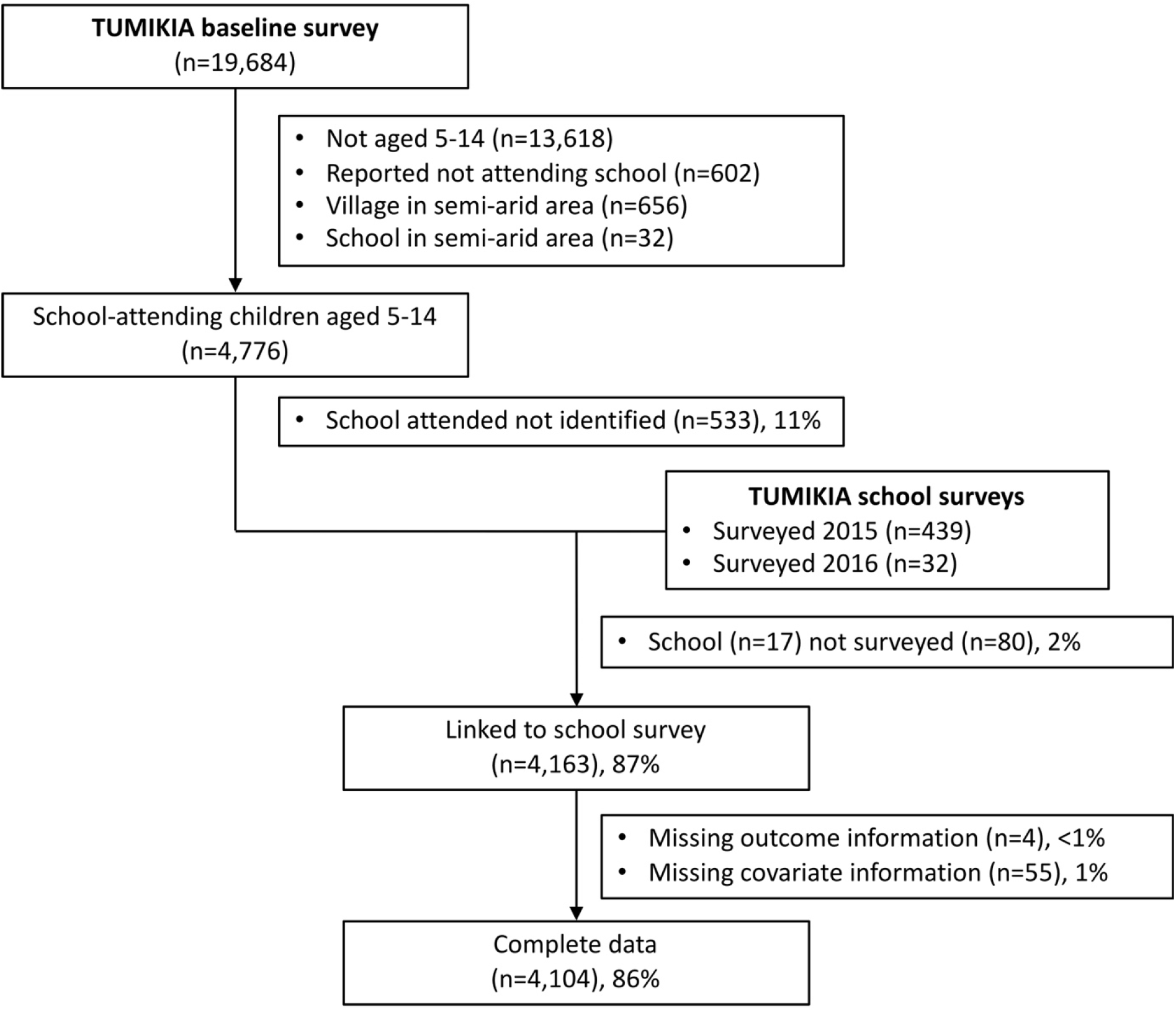
Flow chart of participants in the TUMIKIA baseline survey who were included in the current analysis. Eligible sample included children aged 5-14 years who reported attending school and not residing in villages or attending schools in semi-arid areas. The proportion of eligible subjects with complete data is 86% (4,104/4,776).

Participants resided in 712 villages (median sampled per village 4, range 1, 38) and reported attending 349 schools (median sampled per school 9, Range 1, 49) (Figure 2). Figure 3 describes the structure of the data. In half of all villages, resident children reported attending the same school (range 1 to 8 schools per village). Included schools enrolled children from 1 up to 13 villages. Table 1 describes individual, household, and village characteristics of included children.

**Figure 2.**
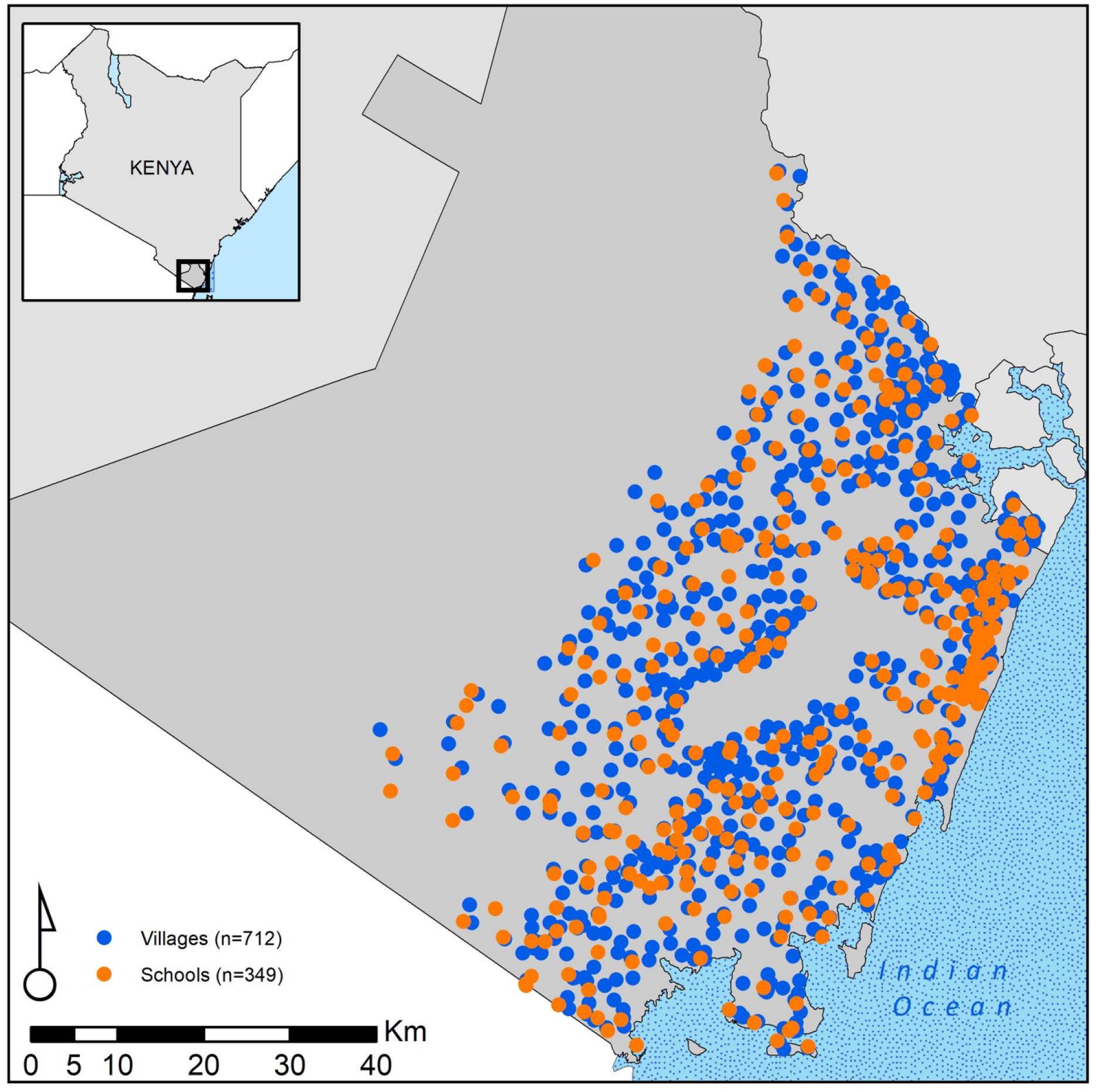
Village of residence and attended schools among 4,104 school-attending children aged 5-14 years in Kwale County, Kenya, 2015.

**Figure 3.**
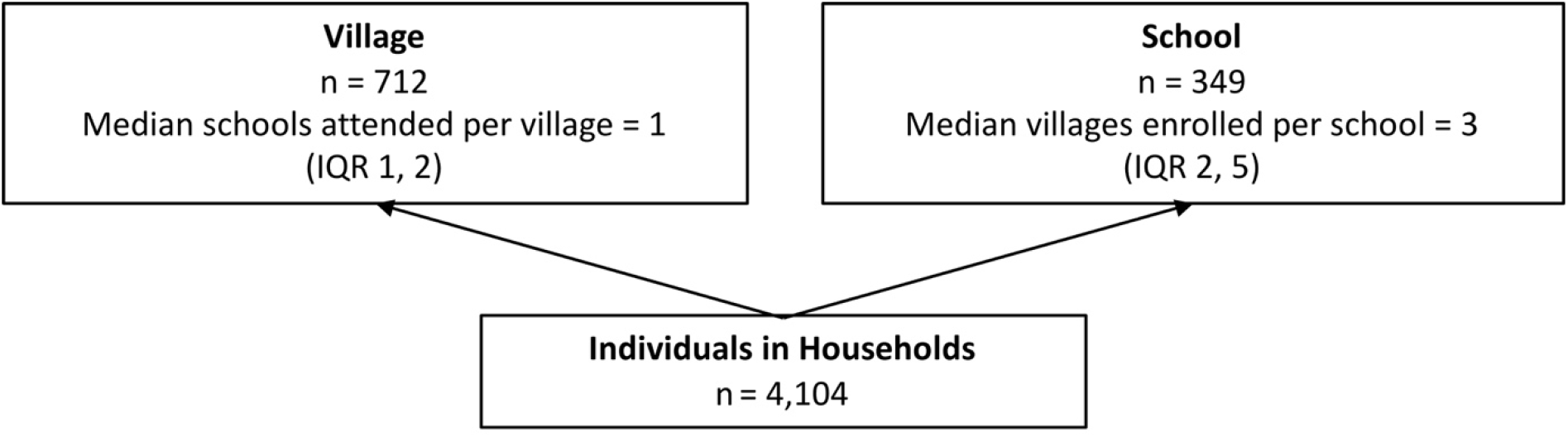
Diagram for the classification model of individuals, villages, and schools.

**Table 1.** Summary characteristics for 4,104 school-attending children in coastal Kenya, 2015

| Characteristics | No. or Mean | % or SD |
| --- | --- | --- |
| **Individual level (n=4104)** |  |  |
| Age (years) | 9.46 | 2.65 |
| Being girls | 2,129 | 51.88 |
| Observed wearing shoes | 1,443 | 35.16 |
| Reported deworming in past year | 2,219 | 54.07 |
| Reported household access to toilet | 2,255 | 54.95 |
| Reported improved water source | 2,216 | 54.00 |
| Missing | 13 | 0.32 |
| Time to fetch water, < 30 min | 3,344 | 81.48 |
| Missing | 14 | 0.34 |
| Covered floor | 833 | 20.30 |
| Household wealth quintile |  |  |
| Most poor, 1 | 1,190 | 29.00 |
| 2 | 2,158 | 52.58 |
| Least poor, 3 | 756 | 18.42 |
| Distance to school (km) (n=3906) | 2.25 | 5.82 |
| **Village level (n=712)** |  |  |
| Proportion reporting toilet access | 0.53 | 0.31 |
| Urbanization |  |  |
| Rural | 545 | 76.54 |
| Periurban | 141 | 19.80 |
| Urban | 26 | 3.65 |
| Aridity index |  |  |
| Dry sub-humid (> 0.5--0.65) | 284 | 39.89 |
| Humid (> 0.65) | 428 | 60.11 |
| Soil sand content $\geq 62\%$ | 256 | 35.96 |
| **School level (n=349)** |  |  |
| Usable toilets per pupil | 0.03 | 0.02 |
| Pupils per usable toilet | 57.79 | 47.35 |
| Total enrolment | 479.47 | 270.67 |
| Urbanization |  |  |
| Rural | 234 | 67.05 |
| Periurban | 84 | 24.07 |
| Urban | 31 | 8.88 |
| Aridity index |  |  |
| Dry sub-humid (> 0.5--0.65) | 130 | 37.25 |
| Humid (> 0.65) | 219 | 62.75 |
| Soil sand content $\geq 62\%$ | 126 | 36.10 |

Overall prevalence of any detected infection was 17.8% and 6.0% for hookworm and *T. trichiura*, respectively, and most infections were classified as light (Table 2). Comparing median odds ratios in tables 3 and 4, school attended had greater impact than village of residence upon hookworm infection (School MOR 2.73, 95% CI 2.50, 2.98; Village MOR 2.09, 95% CI 1.74, 2.38). School and village membership had comparable impacts on *T. trichiura* infection (School MOR 2.92, 95% CI 2.40, 3.43; Village MOR 2.78, 95% CI 2.28, 3.29). Calculated ICC showed similar results. Including measures for sanitation in household, school, and village domains in separate models did not meaningfully change measures of variance and heterogeneity (S5 Table). Including all domain sanitation measures in the same model, we saw no change in calculated MOR and ICC values for hookworm or *T. trichiura* (Tables 3 and 4). Adjusting for potential confounders, residual heterogeneity between schools decreased for *T. trichiura* and, to a lesser extent, hookworm infection, but residual heterogeneity between villages was unchanged.

**Table 2.**
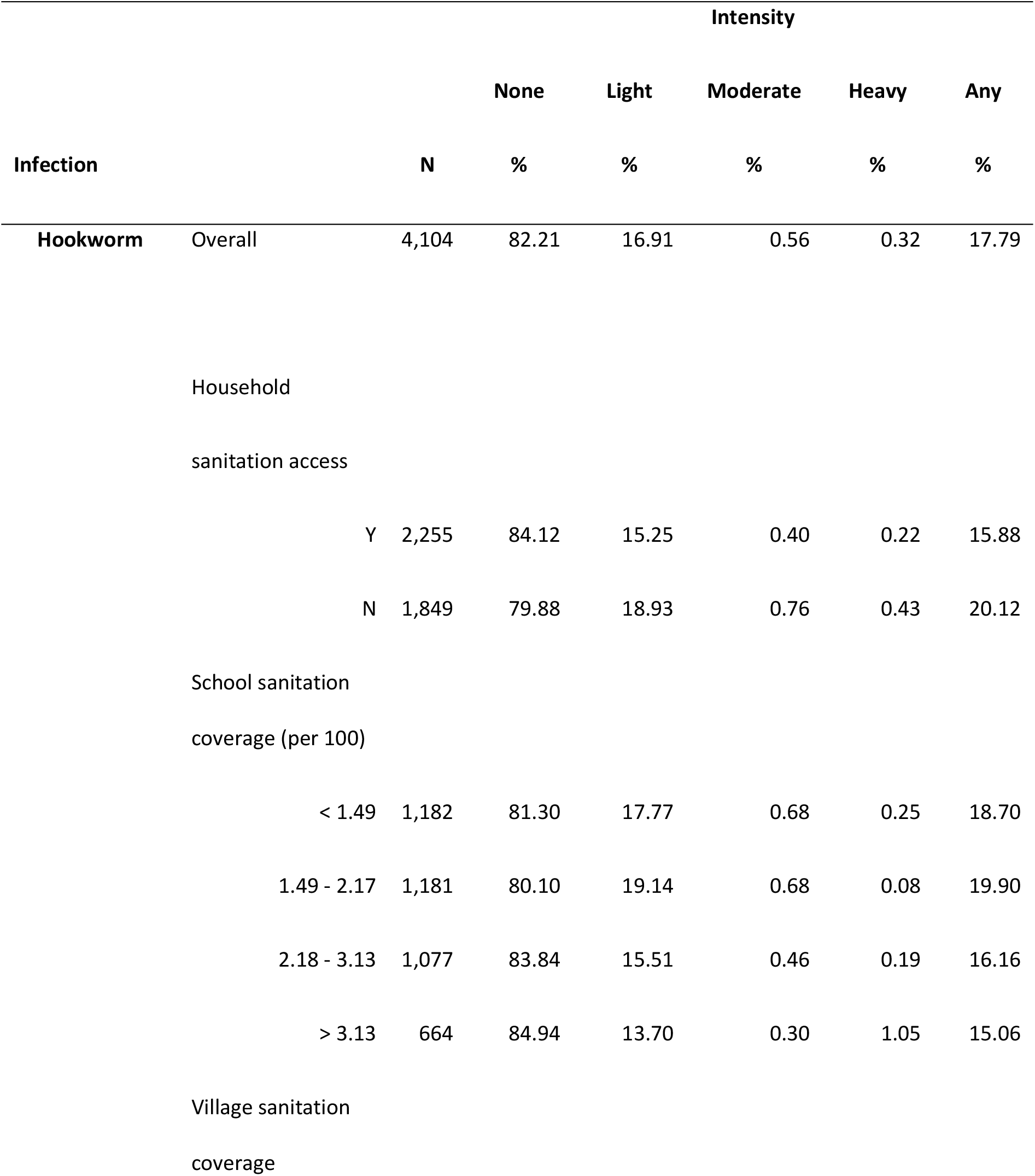
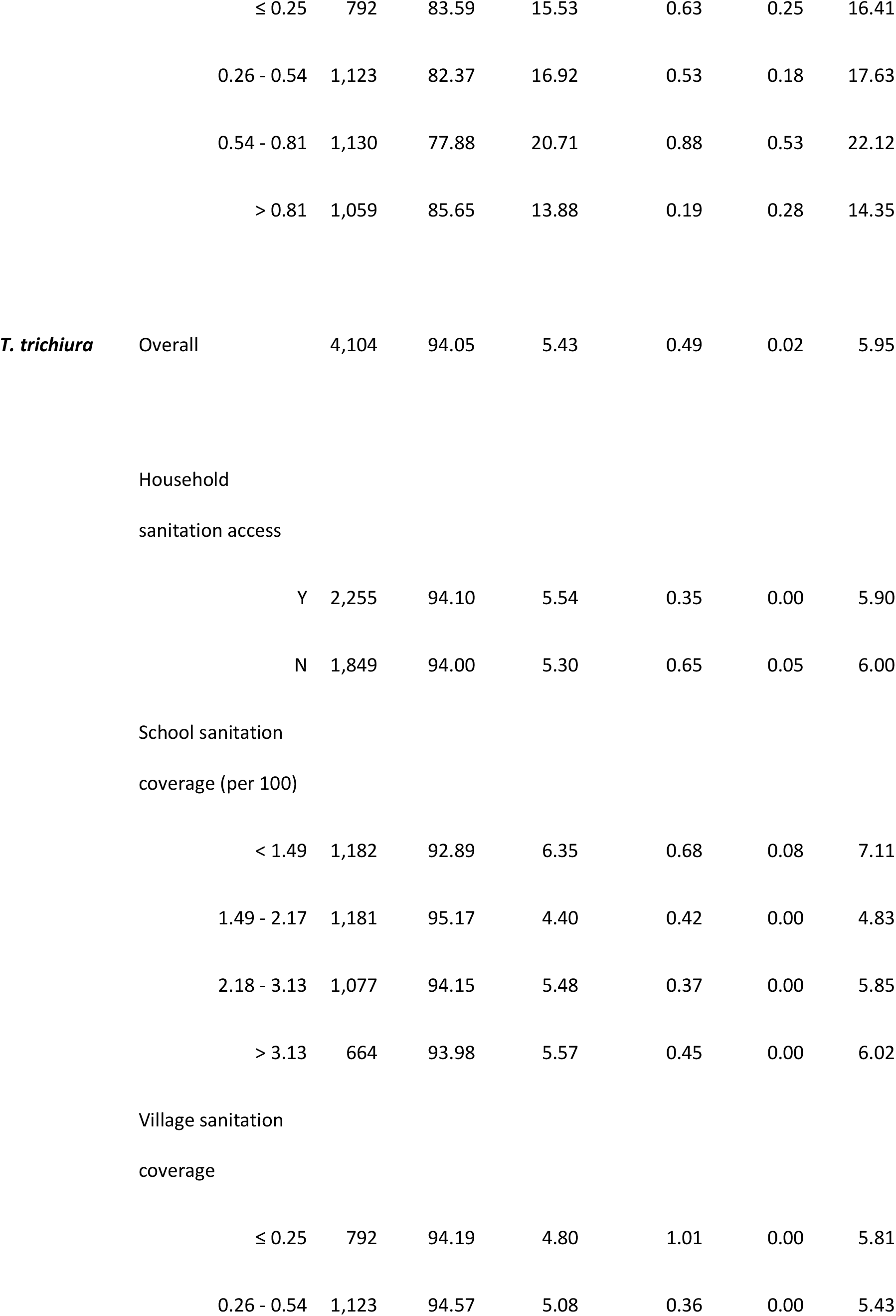
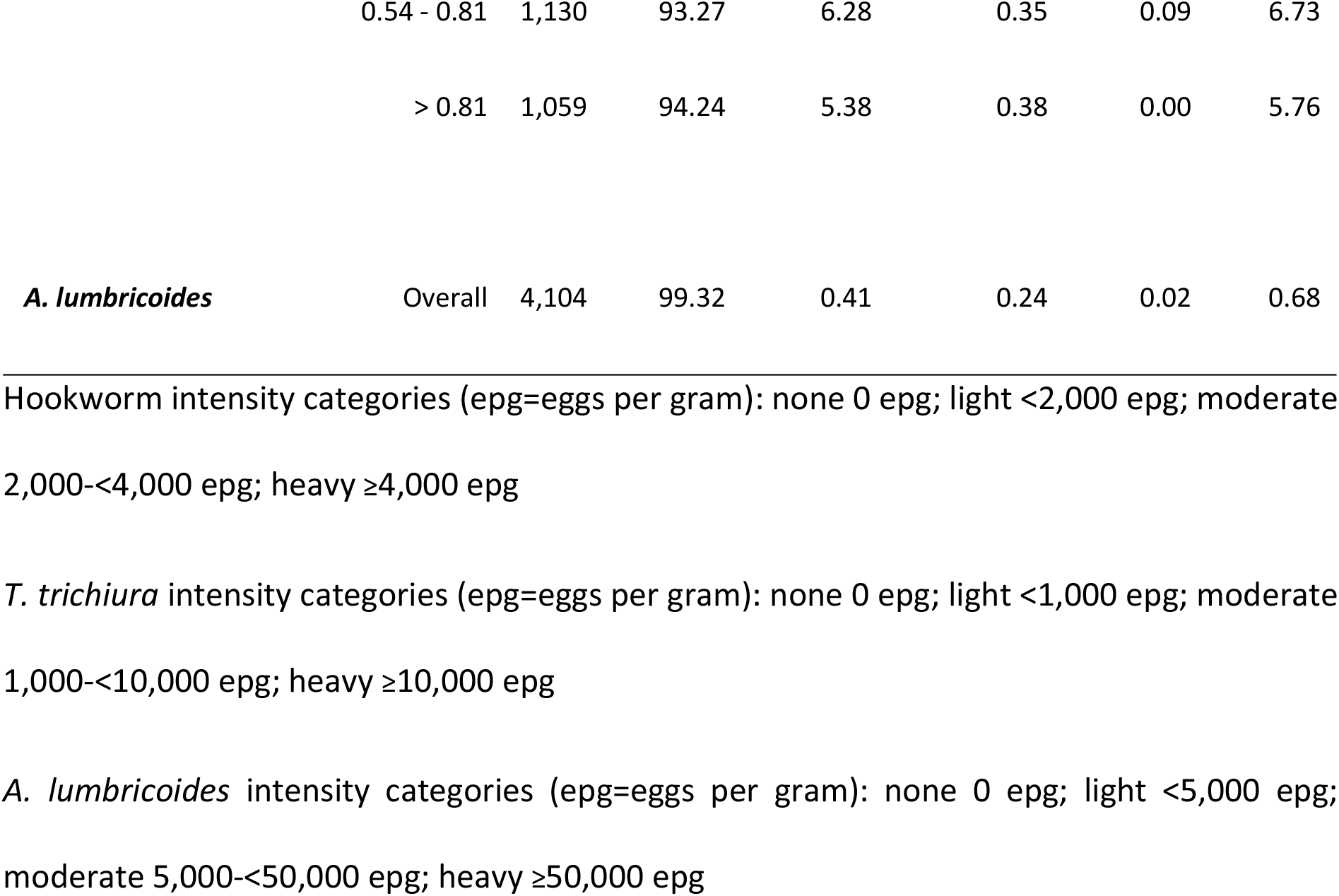
Intensity and any presence of STH infections by household, school, and village sanitation conditions among 4,104 school-attending children in coastal Kenya, 2015.

**Table 3.**
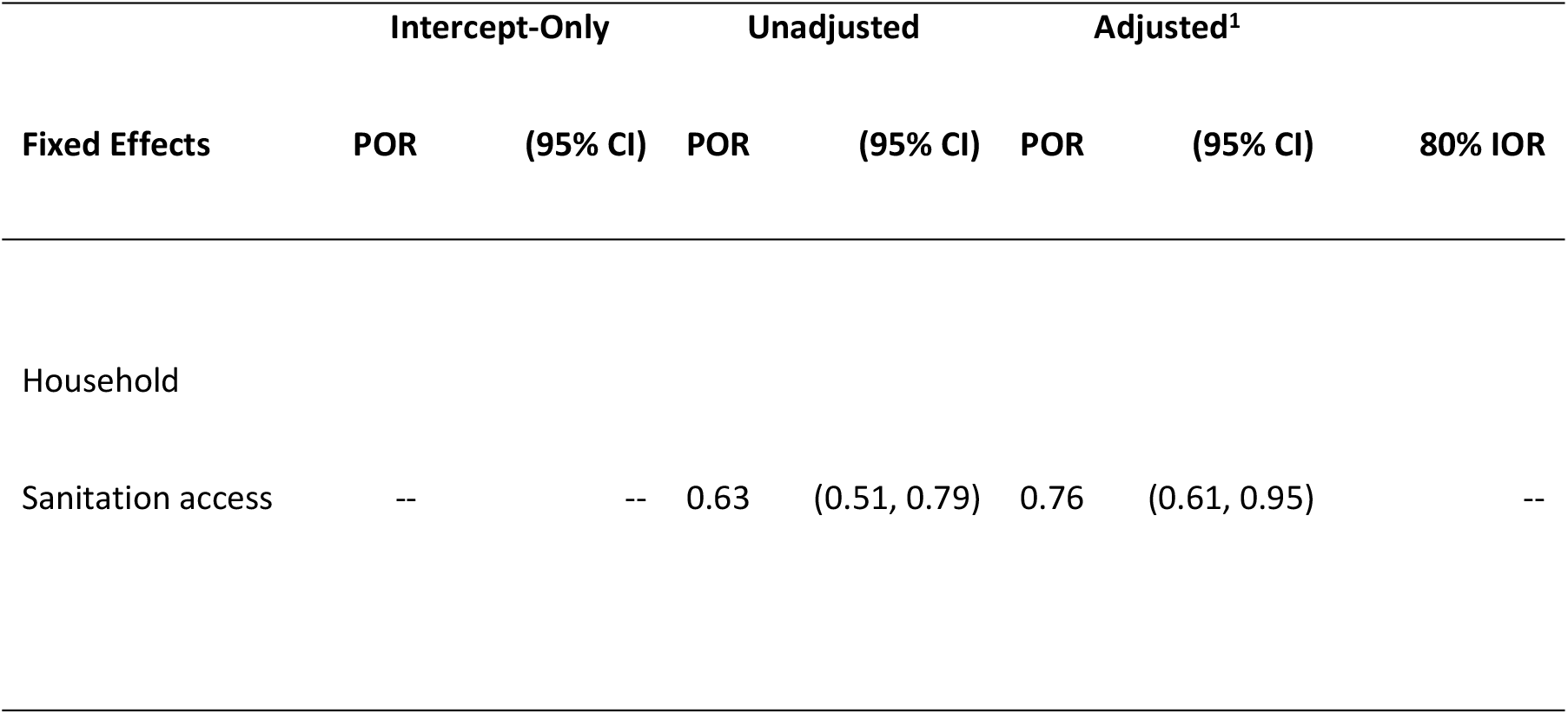
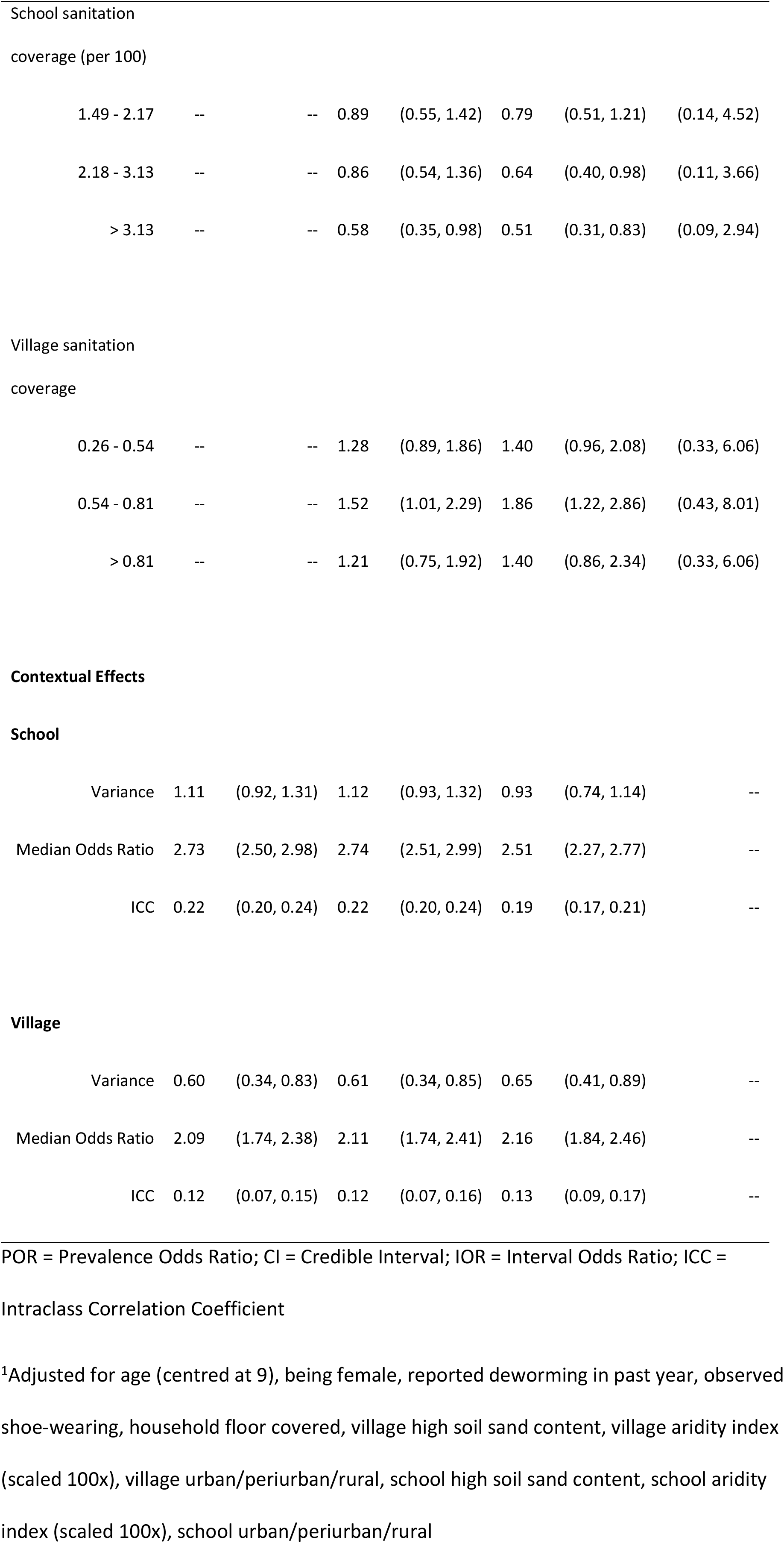
Sanitation and contextual effects on presence of hookworm infection among 4,104 school-attending children in coastal Kenya

|  | Intercept-Only |  | Unadjusted |  | Adjusted<sup>1</sup> |  |  |
| --- | --- | --- | --- | --- | --- | --- | --- |
| Fixed Effects | POR | (95% CI) | POR | (95% CI) | POR | (95% CI) | 80% IOR |
| Household |  |  |  |  |  |  |  |
| Sanitation access | -- | -- | 0.63 | (0.51, 0.79) | 0.76 | (0.61, 0.95) | -- |
| 1.49 - 2.17 | -- | -- | 0.89 | (0.55, 1.42) | 0.79 | (0.51, 1.21) | (0.14, 4.52) |
| 2.18 - 3.13 | -- | -- | 0.86 | (0.54, 1.36) | 0.64 | (0.40, 0.98) | (0.11, 3.66) |
| > 3.13 | -- | -- | 0.58 | (0.35, 0.98) | 0.51 | (0.31, 0.83) | (0.09, 2.94) |
| 0.26 - 0.54 | -- | -- | 1.28 | (0.89, 1.86) | 1.40 | (0.96, 2.08) | (0.33, 6.06) |
| 0.54 - 0.81 | -- | -- | 1.52 | (1.01, 2.29) | 1.86 | (1.22, 2.86) | (0.43, 8.01) |
| > 0.81 | -- | -- | 1.21 | (0.75, 1.92) | 1.40 | (0.86, 2.34) | (0.33, 6.06) |
| **Contextual Effects****School** |  |  |  |  |  |  |  |
| Variance | 1.11 | (0.92, 1.31) | 1.12 | (0.93, 1.32) | 0.93 | (0.74, 1.14) | -- |
| Median Odds Ratio | 2.73 | (2.50, 2.98) | 2.74 | (2.51, 2.99) | 2.51 | (2.27, 2.77) | -- |
| ICC | 0.22 | (0.20, 0.24) | 0.22 | (0.20, 0.24) | 0.19 | (0.17, 0.21) | -- |
| **Village** |  |  |  |  |  |  |  |
| Variance | 0.60 | (0.34, 0.83) | 0.61 | (0.34, 0.85) | 0.65 | (0.41, 0.89) | -- |
| Median Odds Ratio | 2.09 | (1.74, 2.38) | 2.11 | (1.74, 2.41) | 2.16 | (1.84, 2.46) | -- |
| ICC | 0.12 | (0.07, 0.15) | 0.12 | (0.07, 0.16) | 0.13 | (0.09, 0.17) | -- |

POR = Prevalence Odds Ratio; CI = Credible Interval; IOR = Interval Odds Ratio; ICC =
Intraclass Correlation Coefficient
<sup>1</sup>Adjusted for age (centred at 9), being female, reported deworming in past year, observed shoe-wearing, household floor covered, village high soil sand content, village aridity index (scaled 100x), village urban/periurban/rural, school high soil sand content, school aridity index (scaled 100x), school urban/periurban/rural

**Table 4.**
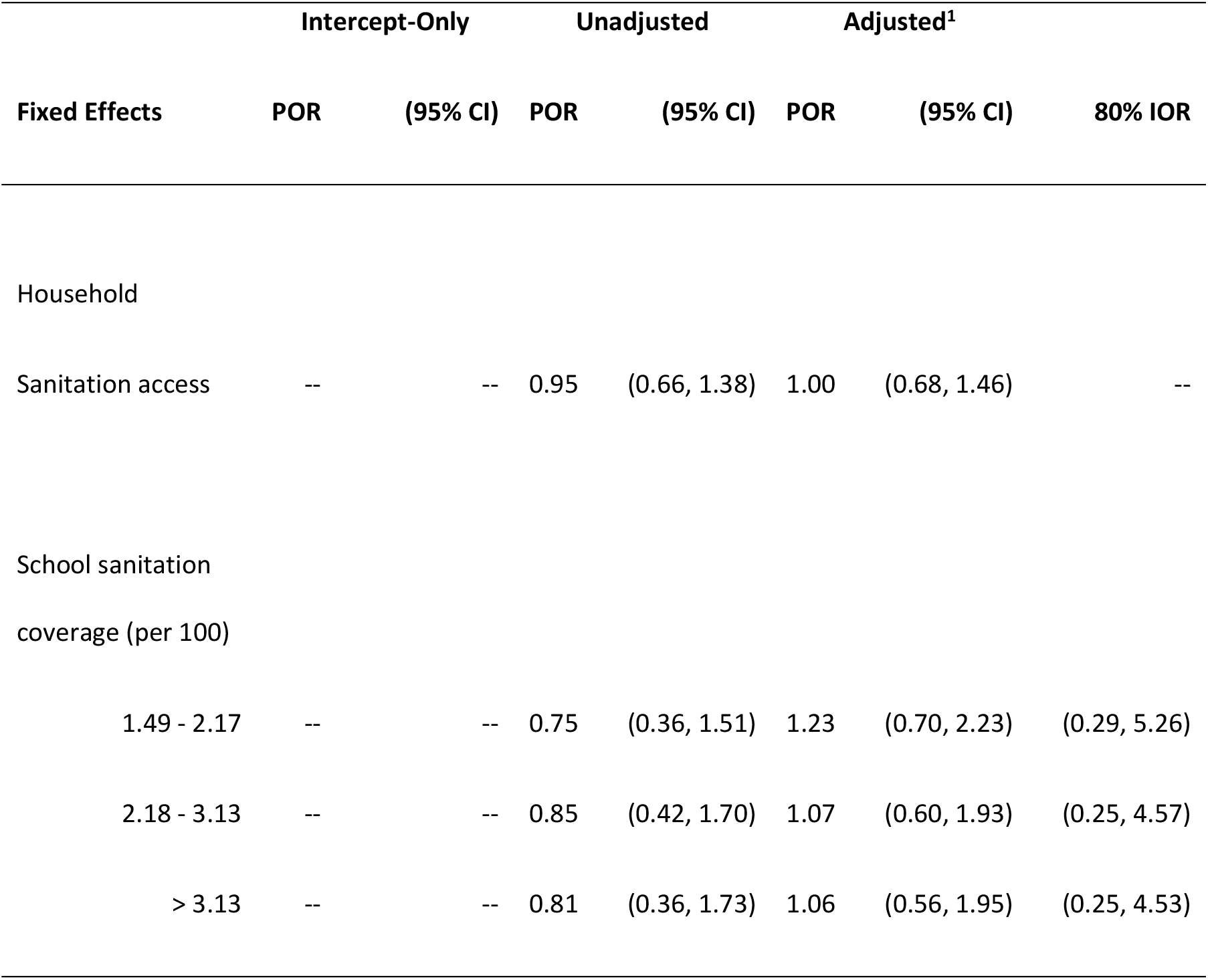
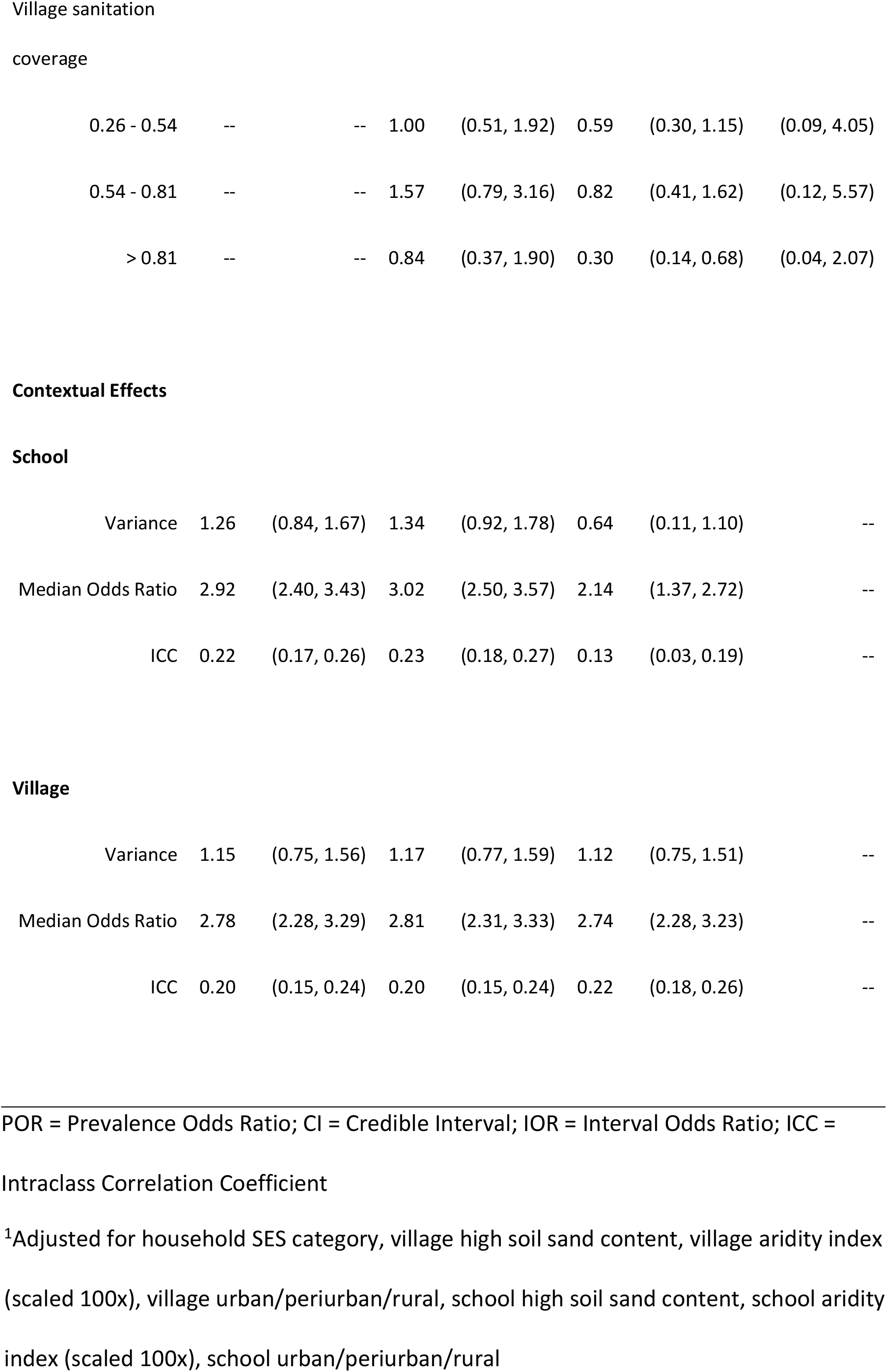
Sanitation and contextual effects on presence of *T. trichiura* infection among 4,104 school-attending children in coastal Kenya

| Fixed Effects | Intercept-Only |  | Unadjusted |  | Adjusted<sup>1</sup> |  |  |
| --- | --- | --- | --- | --- | --- | --- | --- |
|  | POR | (95% CI) | POR | (95% CI) | POR | (95% CI) | 80% IOR |
| Household |  |  |  |  |  |  |  |
| Sanitation access | -- | -- | 0.95 | (0.66, 1.38) | 1.00 | (0.68, 1.46) | -- |
| School sanitation coverage (per 100) |  |  |  |  |  |  |  |
| 1.49 - 2.17 | -- | -- | 0.75 | (0.36, 1.51) | 1.23 | (0.70, 2.23) | (0.29, 5.26) |
| 2.18 - 3.13 | -- | -- | 0.85 | (0.42, 1.70) | 1.07 | (0.60, 1.93) | (0.25, 4.57) |
| > 3.13 | -- | -- | 0.81 | (0.36, 1.73) | 1.06 | (0.56, 1.95) | (0.25, 4.53) |
| coverage |  |  |  |  |  |  |  |
| 0.26 - 0.54 | -- | -- | 1.00 | (0.51, 1.92) | 0.59 | (0.30, 1.15) | (0.09, 4.05) |
| 0.54 - 0.81 | -- | -- | 1.57 | (0.79, 3.16) | 0.82 | (0.41, 1.62) | (0.12, 5.57) |
| > 0.81 | -- | -- | 0.84 | (0.37, 1.90) | 0.30 | (0.14, 0.68) | (0.04, 2.07) |
| School |  |  |  |  |  |  |  |
| Variance | 1.26 | (0.84, 1.67) | 1.34 | (0.92, 1.78) | 0.64 | (0.11, 1.10) | -- |
| Median Odds Ratio | 2.92 | (2.40, 3.43) | 3.02 | (2.50, 3.57) | 2.14 | (1.37, 2.72) | -- |
| ICC | 0.22 | (0.17, 0.26) | 0.23 | (0.18, 0.27) | 0.13 | (0.03, 0.19) | -- |
| Village |  |  |  |  |  |  |  |
| Variance | 1.15 | (0.75, 1.56) | 1.17 | (0.77, 1.59) | 1.12 | (0.75, 1.51) | -- |
| Median Odds Ratio | 2.78 | (2.28, 3.29) | 2.81 | (2.31, 3.33) | 2.74 | (2.28, 3.23) | -- |
| ICC | 0.20 | (0.15, 0.24) | 0.20 | (0.15, 0.24) | 0.22 | (0.18, 0.26) | -- |

POR = Prevalence Odds Ratio; CI = Credible Interval; IOR = Interval Odds Ratio; ICC =
Intraclass Correlation Coefficient
<sup>1</sup>Adjusted for household SES category, village high soil sand content, village aridity index (scaled 100x), village urban/periurban/rural, school high soil sand content, school aridity index (scaled 100x), school urban/periurban/rural

Reported presence of household sanitation access reduced odds of hookworm infection by 37%, compared to no household sanitation access (POR 0.63, 95% CI 0.51, 0.79), among children in villages and schools with similar sanitation conditions. Adjusting for potential confounders, this association was attenuated towards null (POR 0.76, 95% CI 0.61, 0.95). School sanitation coverage in the two highest quartiles (> 2.17 toilets per 100 students) was associated with lower hookworm prevalence, compared to the lowest coverage quartile (< 1.49 toilets per 100 students), adjusting for potential confounders and household and village sanitation (POR 0.64, 95% CI 0.40, 0.98; POR 0.51, 95% CI 0.31, 0.83). No evidence of association of household or school sanitation with *T. trichiura* infection was detected. Children in villages where 54 to 81% of households had sanitation access, compared to children in villages with ≤ 25% sanitation coverage, had 1.86 times the odds of hookworm infection (POR 1.86, 95% CI 1.22, 2.86). Children in villages where > 81% of households had sanitation access, compared to children in villages with ≤ 25% sanitation coverage, had 70% lower odds of *T. trichiura* infection (POR 0.30, 95% CI 0.14, 0.68). Results were robust to the exclusion of 199 children without household coordinates and 197 reporting attending a school > 6.5 km from their house.

Calculated 80% IORs from adjusted models for hookworm and *T. trichiura* were wide and included one for both village and school sanitation measures, indicating sanitation was less important for explaining individual infection, compared to residual variation between these domain levels. For example, comparing children with identical characteristics but drawn from either a village with sanitation coverage ≤ 25% or a village with sanitation coverage > 81%, odds of *T. trichiura* infection will be between 0.04 and 2.07 in 80% of such comparisons.

## Discussion

In this analysis of baseline data from the TUMIKIA trial, we found notable differences between two species of STH in the relationship between sanitation availability and prevalence of infection within different domains among school-attending children in coastal Kenya. Reported use of a sanitation facility by households was associated with reduced prevalence of hookworm infection but was not associated with reduced prevalence of *T. trichiura* infection. Meanwhile, village sanitation coverage > 81% was associated with reduced prevalence of *T. trichiura* infection, but no protective association was detected for hookworm infection. School sanitation coverage > 3.13 toilets per 100 pupils was associated with lower prevalence of hookworm infection. This coverage level, corresponding to a pupil:toilet ratio of 32:1, supports the minimum ratios (25:1 for girls and 35:1 for boys) currently recommended by the Kenyan government (26). School sanitation was not associated with *T. trichiura* infection, however. We found that general contextual effects, represented by residual heterogeneity between village and school domains, had comparable impact upon likelihood of hookworm and *T. trichiura* infection as sanitation coverage in either of these domains.

Three published meta-analyses record considerable heterogeneity in estimates of the effect of sanitation access on hookworm and *T. trichiura* infection. Ziegelbauer et al. found that sanitation was protective against both hookworm and *T. trichiura* infection (5). Strunz et al. found no association of sanitation access with hookworm infection, but it was protective against *T. trichiura* (4). Freeman et al., in the most recent review, found no association of sanitation access with *T. trichiura* infection but sanitation access was protective against hookworm (6). Our findings for household and school sanitation are consistent with these latter results, while adjusting for village and school sanitation coverage plus potential confounders and conditional on village and school membership. Freeman et al. concluded that the lack of an association of sanitation access with *T. trichiura* may result from sanitation’s expected long-term impact on infection compared to a shorter-term impact on reinfection (6). Our findings would support this conclusion. Albendazole, the medication used for Kenya’s National School-Based Deworming Programme (NSBDP), is less effective against *T. trichiura*, compared to hookworm and *A. lumbricoides* (27). Observed associations of household and school sanitation with lower hookworm prevalence in this population of school-attending children could reflect impacts of sanitation in these domains on reinfection, following school-based deworming, rather than a reduction in infection, which might only be observable for *T. trichiura* over a longer time period.

The meta-analyses described above did not distinguish between sanitation access at home or at school. We estimated the independent effects of sanitation access at both the household and school on STH infection. We also expanded upon previous work to estimate the effect of village sanitation coverage (9, 10). The protective association of village sanitation coverage against *T. trichiura* may reflect longer-term impacts of sanitation and a possible community-wide (herd) effect at access levels > 80%, independent of household sanitation access and other factors. In contrast, we found no consistent pattern between village sanitation coverage and hookworm infection. We may not have observed an association because many of the included villages with the lowest sanitation coverage were also in the most arid environments, limiting their suitability for transmission.

Our results clearly show the large contextual effects of village and school domains relative to the estimated fixed effects. The 80% IOR does not indicate precision but provides an interval around our estimated village and school sanitation effects that incorporates unexplained variability between these domains. This result, coupled with the calculated ICCs and MORs, indicates that sanitation coverage in these domains is not a strong predictor of hookworm or *T. trichiura* infection in this setting, though some protective associations were observed. Others have also reported that village membership alone has a large impact on the likelihood of hookworm or *T. trichiura* infection (28), and that heterogeneity of prevalence is associated with multiple environmental and socioeconomic factors (29). Adjusting for potential individual, village and school level confounders in our models did not meaningfully explain further heterogeneity in hookworm infection between villages or schools but did explain some heterogeneity in *T. trichiura* infection between schools.

Though our general contextual effects indicate that village is a relevant context for analysis, the representativeness of village measures is a limitation of the current study. We aggregated village measures from all households sampled for the baseline survey. Because sampling for TUMIKIA was based on CUs, the number of units sampled per village varied. We assumed that included units were representative, but our sample may not have adequately characterised village conditions. Village membership was based on an administrative rather than geographic grouping, so it may also not represent sanitation conditions in the area surrounding study households. While useful for implementation purposes, village may not be the most suitable scale at which to assess community-wide sanitation effects. Future studies could use varying spatial buffers with complete household samples to examine community-wide effects of sanitation coverage on STH infection and identify target thresholds (30). Our household sanitation measure was based on a reported measure and may not reflect actual consistent usage and faeces disposal by household members or faecal contamination levels in the area. Our outcome measure was based on a single stool sample, which may also have underestimated prevalence (31).

In the current study, sanitation conditions, as measured, explained little of the heterogeneity in transmission between villages and schools. Further studies should examine the role of sanitation in different domains against STH infections within the context of school- and community-based mass drug administration. We found evidence of a protective effect of sanitation access at the household against hookworm infection and a sanitation coverage threshold at which a community-wide effect against *T. trichiura* was observed. We also found evidence in support of current school sanitation coverage guidelines towards the control of hookworm infection. In summary, our findings highlight the need for continued efforts, alongside mass drug administration, to extend access to good sanitation facilities at homes, schools, and across communities.

## Data sharing statement

Individual participant data (after de-identification) that underlie the results reported in this paper will be made available on LSHTM Data Compass. The data will be available at the time of publication and researchers can request access through the Data Compass portal. Requests for release of the data will be reviewed by the relevant institutional review boards.

## Acknowledgments

We would like to thank the study participants for their time and patience answering our many questions. We also thank the members of our study team, including supervisors, field officers, laboratory technicians, drivers, and data entry personnel. We thank Claudio Fronterre for advice with modelling and Emma Beaumont for assistance during database assembly and Jessie Hamon for administrative support.

## Supporting Information Legends

**S1 Checklist. STROBE Checklist**

**S2 Supporting Information. Sanitation and hookworm infection model code (dagitty.net)**

**S3 Supporting Information. Sanitation and *Trichuris trichiura* infection model code (dagitty.net)**

**S4 Table. Characteristics of school-attending children in non-arid areas by linked to school survey information status**

**S5 Table. Unadjusted sanitation exposures and contextual effects on presence of hookworm and *T. trichiura* infection among 4104 school-attending children in coastal Kenya**

